# GLOBAL PATTERNS OF PLUMAGE COLOUR EVOLUTION IN ISLAND-LIVING PASSERIFORM BIRDS

**DOI:** 10.1101/2021.09.20.461154

**Authors:** Sean M. Mahoney, Madison D. Oud, Claudie Pageau, Marcio Argollo de Menezes, Nathan Smith, James V. Briskie, Matthew W. Reudink

**Affiliations:** Department of Biological Sciences, Thompson Rivers University, Kamloops, BC, Canada; Physics Institute, Fluminense Federal University, Niteroi, Brazil; National Institute of Science and Technology on Complex Systems, Rio de Janeiro, Brazil; School of Biological Sciences, University of Canterbury, Christchurch, New Zealand

**Keywords:** plumage colour, Passeriformes, ecology and evolution, islands, comparative phylogenetics

## Abstract

Plumage coloration is an important trait involved communication and is shaped by a variety of ecological pressures. Island residency has the potential to change the evolutionary trajectory of plumage colour by differences in habitat and resources, or by altering predation pressure and social selection intensity. Latitude, island size, and isolation may further influence colour evolution by biasing colonization. Therefore, general patterns of plumage evolution are difficult to disentangle. We used phylogenetically controlled analyses to assess the influence of island residency on plumage colouration, by calculating chromaticity values from red, blue, green scores extracted from photos of Order Passeriformes birds. Importantly, we controlled for ecological factors hypothesized to influence colour evolution and assessed family-level effects. We found 1) colour varied between islands and mainlands in females, but not males, and both sexes were affected by several ecological factors; 2) patterns of colour evolution varied among families; 3) island size and distance to the mainland and other islands significantly influenced colour; and 4) interactions between ecological factors and latitude were consistently influenced colour, supporting a latitudinal gradient hypothesis. Our results indicate although island residency influences female colour evolution, a myriad of ecological factors drive plumage colour and the patterns vary among families.

## INTRODUCTION

Animal colour is an important and complex signal used in both inter- and intraspecific interactions (1) and is thought to evolve in response to a variety of evolutionary mechanisms including natural selection, sexual selection, genetic drift, environmental conditions, arbitrary mate choice (2), or some combination of these factors (3). In birds, plumage coloration varies widely among species (4), and colour signals may play important evolutionary roles by mediating mate choice, species recognition, and predator avoidance (5). As such, there is considerable interest in understanding the factors driving global patterns of plumage colour (6), (7),(8),(9),(10),(11).

Island systems provide unique opportunities to explore patterns of plumage colouration. Compared to continental habitats, islands are isolated, relatively small, and are replicated across a broad geographic scale, making them ideal systems to study the evolutionary processes that shape variation in traits (12). In response to insular environments, a number of “island syndrome” studies have documented the parallel evolution of island vertebrates when compared to mainland populations (13),(14). According to the “island rule” (15), body sizes in large vertebrates trend towards dwarfism while small vertebrates trend towards gigantism when comparing island populations to mainland populations (16),(14),(17),(18). Relative to their mainland counterparts, island vertebrates also exhibit K-selected life history strategies, as evidenced by low fecundity, relatively longer developmental periods, and high survival (16), (19). However, island studies have mostly focused on body size and life history traits, while the impact of island environments on ornamental traits is less understood.

Analyses of plumage colouration reveal a general pattern of dull-colouration in island birds, but most of these studies are constrained by small geographic scope and use of relatively few species (7),(8),(10), but see (6). However, a recent worldwide analysis (11) compared plumage colouration of 116 island species to closely related mainland species and found a reduction in plumage brightness and colour intensity as well as a reduction in the number of colour patches in island species. Another large-scale study (20) found colour differences between mainland and island birds in 731 species across three families (Meliphagidae, Fringillidae, and Monarchidae), but the direction of this effect was complex and varied by family: Meliphagidae shifted towards melanin-based plumage while Fringillidae shifted away from carotenoid plumage on island environments. Together, these results suggest different selective pressures may be operating in different lineages or vary geographically.

Several hypotheses have been proposed to explain colour loss in island birds. If plumage colour functions in interspecific interactions, island birds may be duller coloured due to reduced selection for species recognition as island systems typically contain fewer sympatric species than continental areas (7),(8). Alternatively, if exaggerated color expression is under condition dependent sexual selection (21), then island species may become less colourful because of reduced sexual selection pressure on islands (8),(22). Sexual selection is predicted to be relaxed on islands because of reduced genetic diversity from founder effects (23) and/or reduced parasite pressure (24), diminishing the indirect fitness benefits from extra-pair copulations (25). The idea of reduced sexual selection pressure on islands is supported by lower extra-pair paternity rates in island species (26). Changes in the costs of bright plumage may also vary between island and continental habitats. For example, predation pressure on islands is often lower (27) and thus could promote elaboration of plumage colouration rather than camouflage (28),(29). Island species may also show decreased territoriality in part due to fewer con- and heterospecifics, relaxed sexual selection pressure, and/or increased resource availability (11), possibly reducing the need to signal territoriality during species interactions (30). Food resources on islands may differ from those on the mainland, and carotenoid-deficient diets may reduce carotenoid-based (red, orange, yellow) plumage expression (21). Finally, some combination of these effects may result in colour differences between mainland and island birds.

Assessing colour variation in island birds can be challenging due to confounding ecological and natural history factors. First, colour evolution may be affected by latitudinal differences among species. Known as Gloger’s Rule, this biogeographic rule predicts animal coloration will covary with latitudinal changes in body temperature regulation, colours needed for camouflage, parasite loads, or some combination of these factors (31). Second, differences in habitat may influence ambient light in the environment, so there may be selection for or against ornate plumage given habitat-specific light conditions (32). Third, island characteristics such as geographic size and isolation (i.e., distance from mainland) may affect the evolutionary trajectory of colour evolution by biasing colonization or by limiting population sizes, thereby diminishing or exacerbating genetic drift effects (23). Finally, macroevolutionary studies often assess higher-level taxonomic processes, but because selection pressures likely vary among families, the directionality of the effects may change at lower taxonomic levels (33),(34). Therefore, to unravel the mechanisms mediating plumage evolution, it is important for studies to assess the biological and ecological factors contributing to colour at a global scale and across a broad range of taxa.

Although previous studies have documented colour differences between mainland and island bird populations using a few families or subsets of species (e.g., (11),(20)), no study has comprehensively assessed the selective pressures driving plumage colour evolution in island birds using an entire order of birds—while also simultaneously controlling for confounding biotic and abiotic factors—calling into question the generalizability of an island effect on plumage coloration. Using a phylogenetic statistical framework, in this study we leveraged a global and comprehensive dataset of plumage colour in the Order Passeriformes to test the hypothesis that colour would differ between mainland and island populations. The Passeriformes is an ideal order to test hypotheses on plumage colour evolution because 1) it is a speciose order (more than half of all living birds), 2) passeriform species generally exhibit relatively ornate plumage colouration (6), 3) there is high variation in plumage colour among species, and 4) passeriform birds are broadly distributed throughout the world in both mainland and island habitats (35). We specifically tested the prediction that male and female passerines occupying islands would be less colourful than those occupying the mainland (*sensu* (11)), while also controlling for covariates such as latitude (i.e. “Gloger’s Rule”, (31)), diet (3), and variation in ambient light in the habitat (32). Given that selection pressures may vary among species within Passeriformes, we also tested for colour differences between mainland and islands species at the family-level. Finally, because island size and island isolation may influence species richness, resource availability, predation pressure, and/or bias island colonization (*sensu* (11),(29)), we also tested the effect of island size and isolation on plumage coloration.

## MATERIALS AND METHODS

### Data Collection

We classified 5,693 extant Passeriformes species (Dryad Data Respository) as mainland or island dwelling (Fig. 1) using global range maps from the International Union of Conservation of Nature’s Red List of Threatened Species (36). We considered mainland to be a land mass larger or equal to 7.7 million km^2^ (approximately the area of mainland Australia, the smallest defined continent (37)). We defined islands as smaller or equal to 2.2 million km^2^ (approximately the size of Greenland, the largest defined island (37),(38). Passerines where 80% of their range covered non-continental landmasses (such as the Hawaiian islands or New Zealand) were classified as “island” species (n=1183). Species were classified as “mainland” when approximately 80% or more of the range covered a continent (such as North America or Australia) (n=4510). Throughout, we use the term land classification to refer to designation of a species as either an island or continental species Using IUCN classifications, we also collected the latitude centroid of each species range. Diet, habitat type, and geographic region were taken from (39) based on (40), (41), and (42). These factors may influence plumage colouration differences between island and continental species because of variation in pressures associated with thermoregulation, resource availability, dietary precursors, or light environments. We obtained data on island size, distance to nearest mainland, and distance to nearest other islands from the UN Environmental Programme island directory (38). Island size may influence colour evolution because there may be a greater diversity of predators on larger islands (29), while a lower diversity of congeneric species on smaller islands may reduce species recognition pressures (11). Distance to continents and other islands may influence the evolutionary trajectory of colouration of island birds through founder effects of relatively few individuals and alleles (35). Collectively, our dataset included land classification (island vs. continental), habitat, diet, global geographic region, range latitude for all extant passerines, and island size, distance to mainland, and distance to other islands for passerines found on islands (Dryad Data Repository).

**Fig. 1.**
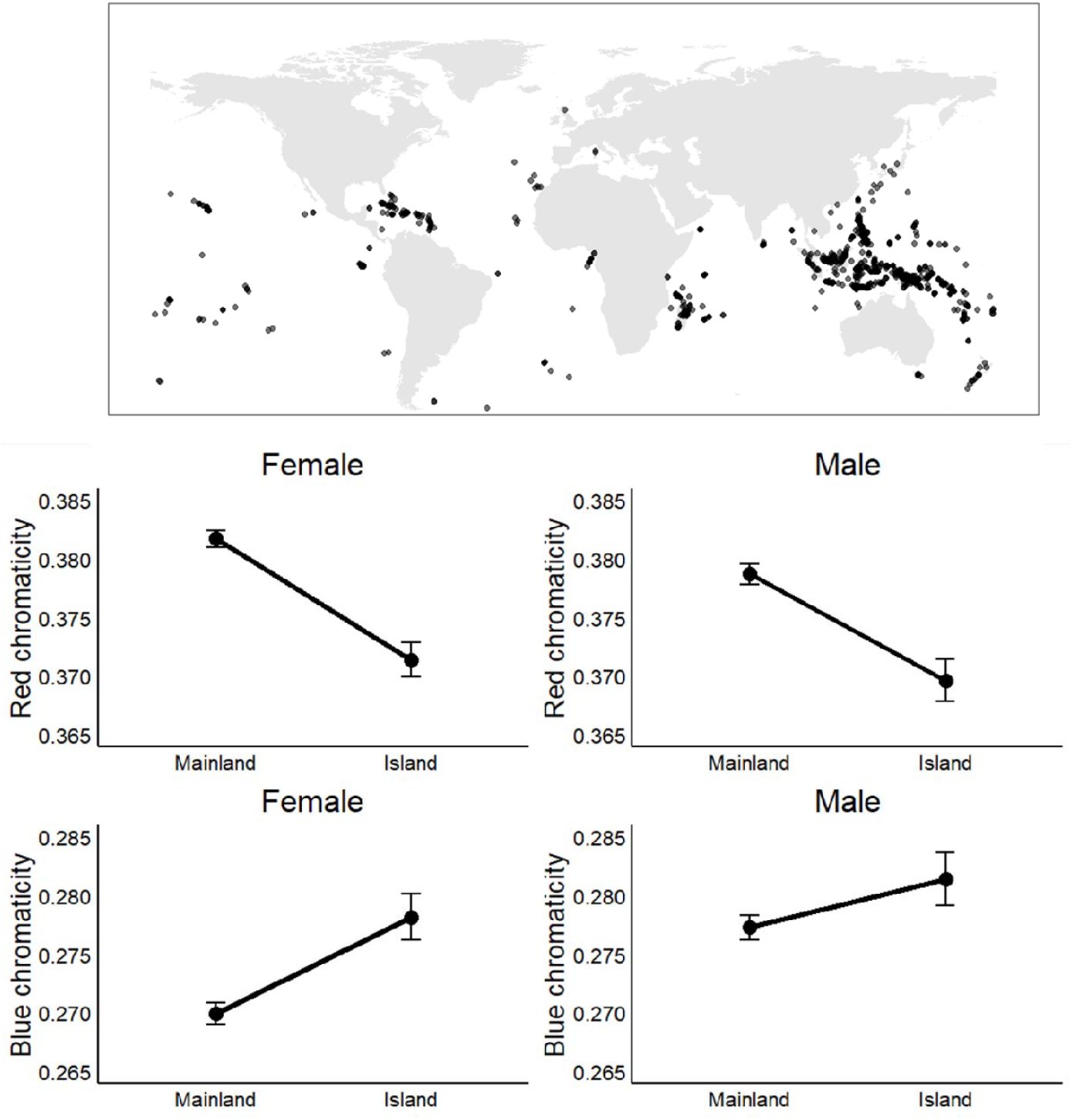
Top panels: Global distribution of Passeriformes island species (n=1,183) used in the present study. Bottom panels: Female and male colour variation between mainland and island passerine birds (n=5,693). Red chromaticity (±SE) was significantly higher on the mainland for females (F=57.2, P<0.0001) but not in males (F=0.05, P=0.82). Blue chromaticity (±SE) was significantly lower on the mainland for females (F=23.7, P<0.0001) but did not differ in males (F=2.2, P=0.14).

We used plumage colour data from (43), who quantified red, green, and blue (RGB) values from images of the crown, forehead, nape, throat, upper breast and lower breast of all the Passeriformes species listed in the Birds of the World (44). Because carotenoid- and structural-based colouration lead to elaborate colouration through different physiological mechanisms, we used colour scores extracted from the RGB values (43) to calculate chromaticity. Red chromaticity was calculated using the equation: R/(R+G+B), and blue chromaticity was calculated using the equation: B/(R+G+B), where R is the red value, G is the green value, and B is the blue value from (43). Chromaticity provides an estimate of the relative short and long wavelengths and is a reliable estimate of carotenoid and structural plumage coloration (45).

To assess the efficacy of using chromaticity to estimate “carotenoid” and “structural” plumage coloration, a single, independent observer categorized the colour for multiple plumage patches in males and females of all species in the Family Thraupidae (n=346 species/sex) using the publicly available visual media source Birds of the World by the Cornell Lab of Ornithology (44). In this analysis, we used 10 patches: auricular, crown, back, rump, throat, breast, belly, crissum, wingtip (color of the tip of the longest primary flight feather), and dorsal side of the rectrices. Patches were chosen to mirror (46), who measured the reflectance spectrum of each patch colour using a spectrophotometer for most members in Thraupidae. We then compared the chromaticity values to these classifications and found the distribution of red and blue chromaticity scores did not overlap in plumage patches classified as “blue” or “red.” This suggests that our chromaticity values effectively captured the variation in structural blue and carotenoid red plumage coloration (Fig. S1).

### Phylogenetic Methods

To control for phylogenetic relationships in our analyses, we downloaded 1000 potential phylogenies from birdtree.org (47),(48) for the 5,693 passerine species included in the dataset. We used TreeAnnotator in BEAST v1.10.1 (49) to construct a maximum clade credibility tree using 1% burn in and mean node heights. We repeated these steps with the 1,183 island passerines to test the effect of island characteristics on passerine colour.

### Statistical Analysis

We performed all analyses in R 3.5.3 (50) using phylogenetic generalized least squares (PGLS) in the *nlme* package (51). We tested how male and female passerine colour variation was explained by land classification, diet, habitat, latitude, and region using stepwise model reduction based on Akaike Information Criterion (AIC). We first built a full model, which included either red or blue chromaticity as the response variable, and land classification, diet, habitat, latitude, region, and their interactions (land classification x latitude, land classification x diet, land classification x habitat, and habitat x diet) as the main effects. We built separate models for each sex. We then undertook model reduction for all possible models using the *StepAIC* function in the *MASS* package and selected the top model based on the change in AIC (ΔAIC, (52)) between the full model and each reduced model. We considered ΔAIC values within 4 to be competitive and chose our final model based on the lowest AIC (52). To assess differences in directionality among families, we included family as a fixed effect in the final model and plotted the results for each family. We then repeated these steps using only island passerines and included island size, distance to mainland, and distance to other islands and two-way interactions between all terms. We used phylogenetic path analyses using the R package *phylopath* to assess the direct and indirect effects of the variables from the top PGLS models. We first built candidate path analyses informed by the PGLS models and then ranked models using an information theory approach based on C-statistics (53). Information theory evaluates the conditional independencies of each model and assigns a C statistic. The models are ranked based on the change in C statistic (ΔCICc) between models, where lower C scores are optimized models and ΔCICc <2 are competitive. The top phylogenetic path analysis model was then selected as the model with lowest C statistic and <2 ΔCICc.

## RESULTS

### Effect of island residency and ecological factors on Passeriformes colour evolution

We identified the reduced PGLS model as the top models for female red chromaticity (Table S1) that included land classification (F=57.21, P<0.0001), habitat (F=22.40, P<0.0001), diet guild (F=5.69, P=0.003), latitude (F=65.39, P<0.0001), region (F=15.54, P<0.0001), and the interactions between land classification and latitude (F=5.51, P=0.001), land classification and diet guild (F=10.60, P<0.0001), and habitat and diet guild (Table 1, F=4.55, P<0.0001). Similarly, the top model for male red chromaticity was the reduced model (Table S1) and included land classification (F=0.05, P=0.82), habitat (F=1.17, P=0.32), diet guild (F=22.46, P<0.0001), latitude (F=15.81, P<0.0001), region (F=11.28, P<0.0001), and the interactions between land classification and latitude (F=2.17, P=0.14), land classification and diet guild (F=25.06, P<0.0001), and habitat and diet guild (Table 1, F=2.61, P=0.02).

**Table 1.** AIC selected model results demonstrating the effect of each fixed effect and interactions on female and male Passeriformes plumage coloration. Significant results are indicated in bold text.

| Sex | Variable | Fixed effect | *df* | *F* | *P* |
| --- | --- | --- | --- | --- | --- |
| Female | Red chromaticity | **Land classification** | **1** | **57.21** | **<0.0001** |
|  |  | **Habitat** | **3** | **22.40** | **<0.0001** |
|  |  | **Diet** | **2** | **5.69** | **0.003** |
|  |  | **Latitude** | **1** | **65.39** | **<0.0001** |
|  |  | **Geographic region** | **6** | **15.54** | **<0.0001** |
|  |  | **Land x Latitude** | **3** | **5.51** | **0.001** |
|  |  | **Land x diet** | **2** | **10.60** | **<0.0001** |
|  |  | **Habitat x diet** | **6** | **4.55** | **<0.0001** |
|  | Blue chromaticity | **Land classification** | **1** | **23.72** | **<0.0001** |
|  |  | **Habitat** | **3** | **20.09** | **<0.0001** |
|  |  | Diet | 2 | 3.00 | 0.05 |
|  |  | **Latitude** | **1** | **75.58** | **<0.0001** |
|  |  | **Geographic region** | **6** | **7.31** | **<0.0001** |
|  |  | Land x Latitude | 1 | 2.54 | 0.11 |
|  |  | **Land x diet** | **2** | **17.93** | **<0.0001** |
|  |  | **Habitat x diet** | **6** | **3.72** | **<0.0001** |
| Male | Red chromaticity | Land classification | 1 | 0.05 | 0.82 |
|  |  | Habitat | 3 | 1.17 | 0.32 |
|  |  | **Diet** | **2** | **22.46** | **<0.0001** |
|  |  | **Latitude** | **1** | **15.81** | **<0.0001** |
|  |  | **Geographic region** | **6** | **11.28** | **<0.0001** |
|  |  | Land x Latitude | 1 | 2.17 | 0.14 |
|  |  | **Land x diet** | **2** | **25.06** | **<0.0001** |
|  |  | **Habitat x diet** | **6** | **2.61** | **0.02** |
|  | Blue chromaticity | Land classification | 1 | 2.21 | 0.14 |
|  |  | Habitat | 3 | 1.96 | 0.12 |
|  |  | **Diet** | **2** | **30.09** | **<0.0001** |
|  |  | **Latitude** | **1** | **14.01** | **<0.0001** |
|  |  | **Geographic region** | **6** | **7.55** | **<0.0001** |
|  |  | **Land x diet** | **2** | **21.13** | **<0.0001** |
|  |  | **Habitat x diet** | **6** | **2.72** | **0.01** |

The top model for female blue chromaticity was the reduced model (Table S1) and was explained by land classification (F=23.72, P<0.0001), habitat (F=20.09, P<0.0001), diet guild (F=3.0, P=0.05), latitude (F=75.58, P<0.0001), region (F=7.31, P<0.0001), the interactions between land classification and diet guild (F=17.93, P<0.0001), and habitat and diet guild (Table 1, F=3.72, P<0.0001), but not land classification and latitude (F=2.54, P=0.11). Blue chromaticity in males was best explained by the reduced model (Table S1) and included diet guild (F=30.09, P<0.0001), latitude (F=14.01, P<0.0001), region (F=7.55, P<0.0001), and the interactions between land classification and guild (F=21.13, P<0.0001), and habitat and diet guild (Table 1, F=2.72, P=0.01), but not land classification (F=2.21, P=0.14) or habitat (F=1.96, P=0.12).

In general, we found mixed support for our prediction that island birds have reduced coloration. Colour differed significantly between mainland and island females, but not males (Fig. 1). Female red chromaticity was higher on the mainland than on islands (Fig. 1, F=57.21, P<0.0001) while female blue chromaticity was lower on the mainland relative to islands (Fig. 1, F=23.72, P<0.0001).

Apart from the effects of islands on plumage, we found colour varied with several ecological and natural history covariates and their interactions, underscoring the complexity of the macroevolutionary processes driving plumage colour evolution (Table 1). In support of Gloger’s Rule, colour varied by latitude (Fig. S2): red chromaticity was higher and blue chromaticity was lower near the equator for both males (F=15.81, P<0.0001) and females (F=65.39, P<0.0001). However, the interaction between land classification and latitude revealed red chromaticity in island females was positively related to latitude, while mainland red chromaticity was negatively related to latitude (Fig. S2, F=5.51, P=0.001).

Colour also varied with diet, where both invertivore and omnivore guilds had higher red chromaticity (female: F=5.69, P=0.003; male: F=22.46, P<0.0001) and blue chromaticity (Table 1, male: F=30.09, P<0.0001). However, the interactions between diet guild and land classification indicated that red chromaticity in invertivores was lower on islands (female: F=10.60, P<0.0001; male: F=25.06, P<0.0001). Blue chromaticity in invertivores and herbivores was higher on islands but did not vary between islands and mainlands for omnivores (Fig. S3, female: F=17.93, P<0.0001; male: F=21.13, P<0.0001).

Colour varied among habitat types in females, with species in open habitats having higher red chromaticity (F=22.40, P<0.0001) while those in dense habitats had lower blue chromaticity (Table 1, F=20.09, P<0.0001). The interaction between habitat and guild indicates red chromaticity in herbivorous birds was higher in dense and open habitats, but lower in aquatic habitats (F=4.55, P<0.0001). Similarly, blue chromaticity in herbivorous birds was lower in dense and open habitats but higher in aquatic habitats (Table 1, F=3.72, P<0.0001).

### Phylogenetic Path Analyses

Our phylogenetic path analyses indicated that plumage colour in the Passeriformes is influenced by several factors (Table S2). The top models predicting red chromaticity in females were explained by the direct effects of habitat and land classification and the indirect effect of habitat on land classification (Fig. 2, CICc=451.6), while female blue chromaticity was explained by geographic region (Fig. 2, CICc=506.2). The top model predicting red chromaticity in males was explained by the direct effects of land classification and latitude, and the indirect effect of latitude on land classification (Fig. 2, CICc=399.6). Blue chromaticity in males was explained by the direct effect of diet and land classification and the indirect effect of diet on land classification (Fig. 2, CICc=486.9).

**Fig. 2.**
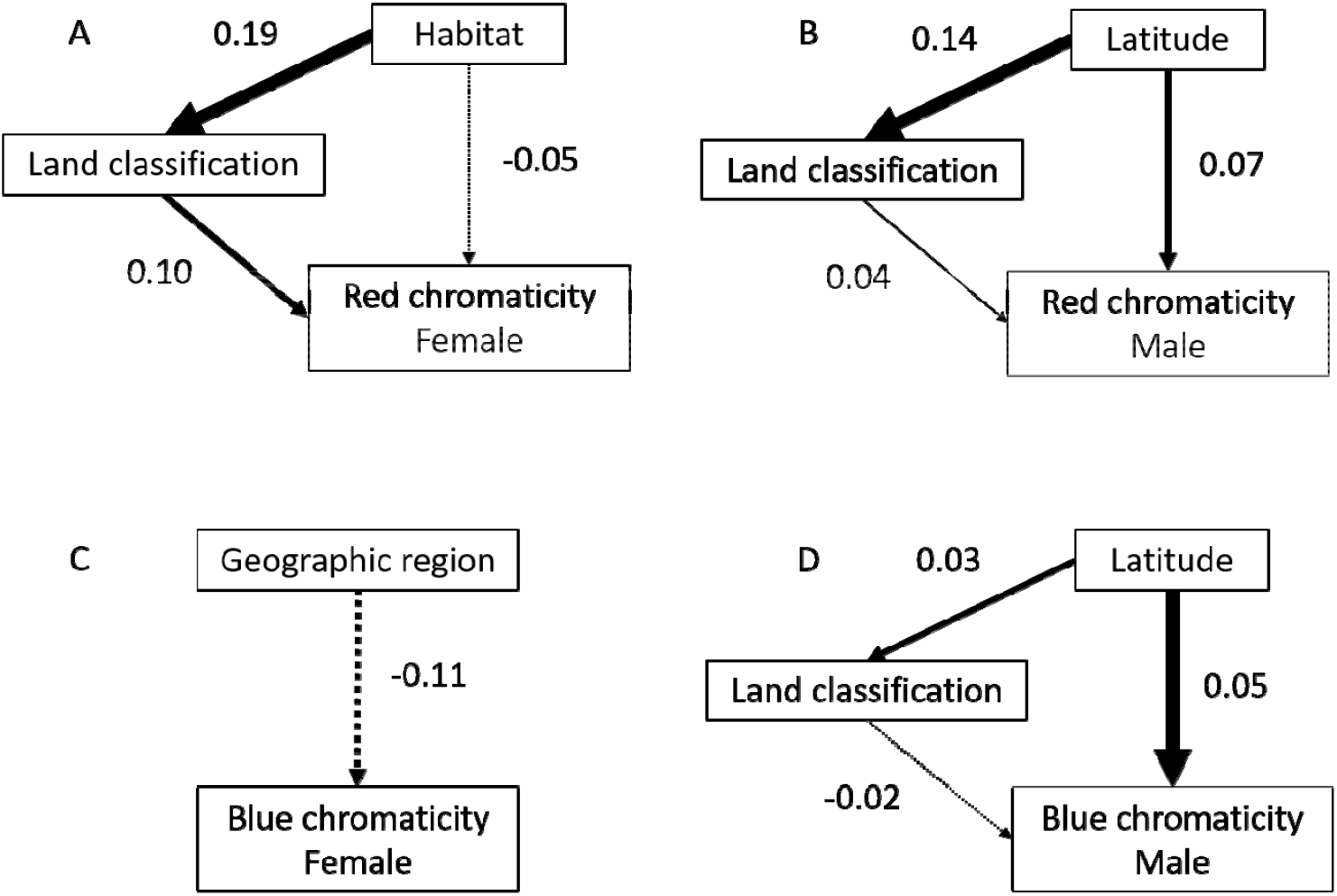
Final path analysis models illustrating the effect of A) habitat and land classification (island/mainland) on female red chromaticity, B) geographic region on blue chromaticity, C) latitude and land classification on male red chromaticity, and D) male blue chromaticity. Solid lines indicate positive, while dashed lines indicate negative effects.

### The Island Effect varies among Passeriformes families

The direction of colour change between mainland and island birds varied among families in the Passeriformes (Fig. 3). Red chromaticity increased in island females and males in Fringillidae, Meliphagidae, and Sturnidae but decreased in Estrildidae, Pellorneidae, Pycnonotidae, Tyrannidae, and Vireonidae (Fig. S4). Additionally, red chromaticity in females in the Parulidae and Turdidae, and in males in the Muscicapidae and Oriolidae decreased on islands (Fig. S4). Blue chromaticity increased in island females and males in Oriolidae, Pellornidae, Pycnonotidae, Turdidae, Tyrannidae, and Vireonidae but decreased in Sturnidae and Zosteropidae (Fig. S4). Blue chromaticity in island females increased in Campephagidae and decreased in Meliphagidae (Fig. S4). Male blue chromaticity increased in Muscicapidae and decreased in Fringillidae and Ploceidae between the mainland and islands (Fig. S4).

**Fig. 3.**
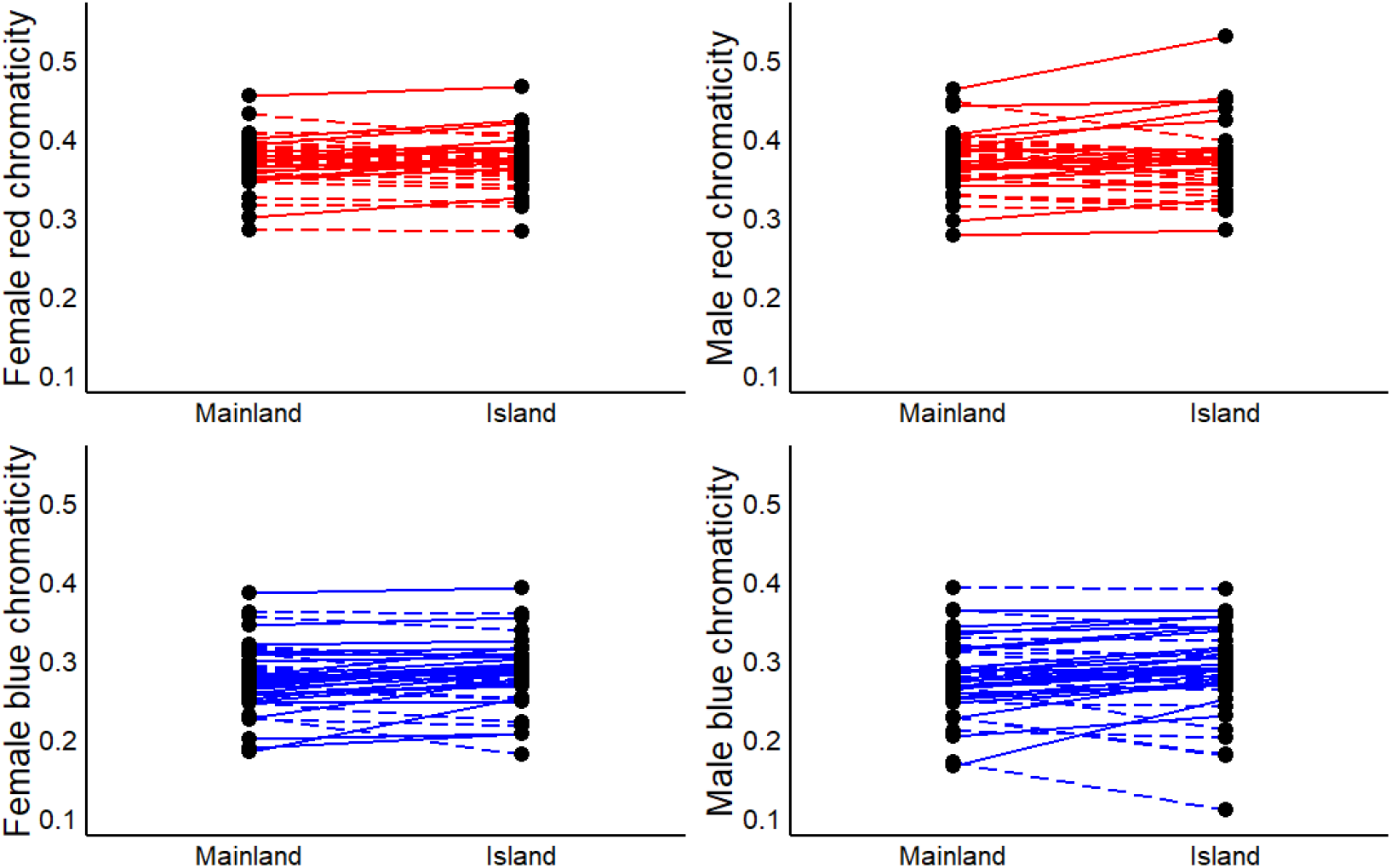
Colour variation between mainland and island species among Passeriformes (dots). Solid lines represent increases in red (top panels) and blue chromaticity (bottom panels) and dashed lines indicate decreases. Although generally female red decreases and blue increases on islands, the direction of this effect varies among families.

### Island characteristics predict colour evolution within island Passeriformes

In our analyses restricted to only island birds, we found the top models for red and blue chromaticity were the reduced models (Table S3) and included island size, island isolation (i.e. distance to the mainland and other islands), and other ecological covariates as above (Table S4). In females, red and blue chromaticity varied with island size, where red was lower and blue was higher on larger islands (Table S4, red: F=17.64, P<0.0001; blue: F=12.33, P<0.0001) and by island isolation (Fig. S5). Whereas female red chromaticity increased on more distant islands (distance to the mainland: F=5.69, P=0.02; distance to other islands: F=25.35, P<0.0001, Fig. S5), blue chromaticity decreased on more isolated islands (Fig. S5, distance to other islands: F=18.84, P<0.0001). We found two-way interactions between latitude and island size and latitude and island isolation, which revealed that red chromaticity increased at higher latitudes regardless of island size (Table S4, F=10.74, P<0.0001), and on more isolated islands, regardless of latitude (Table S4, latitude x nearest mainland: F=8.93, P<0.0001; latitude x nearest islands: F=9.11, P<0.0001). Blue chromaticity decreased on larger islands (Table S4, F=12.33, P<0.0001) and on more isolated islands (Fig. S5, nearest island: F=18.84, P<0.0001). We also found an interaction between island area and island isolation in blue, indicating blue chromaticity was lower on more distant islands regardless of island size (Table S4, nearest island: F=5.24, P=0.02). Further, there was an interaction between latitude and geographic region (Table S4). While birds in most regions showed increases in red chromaticity and decreases in blue chromaticity, birds in Australasia exhibited decreases in red and increases in blue across latitude (Table S4, F=4.96, P<0.0001). Additionally, there was an interaction between land classification and diet in female blue chromaticity, indicating blue decreased across latitude in herbivores and increased in invertivores, but blue did not in ominvores (F=4.21, P=0.02).

Similarly in males, red and blue chromaticity varied by island size, island isolation, and other ecological covariates (Table S3). Male red chromaticity was lower and blue chromaticity was higher on larger islands (Table S4, red: F=18.92, P<0.0001, blue: F=12.37, P<0.0001). In contrast, red was higher, and blue was lower on more isolated islands (Fig. S5, red nearest mainland: F=10.45, P<0.0001; blue nearest mainland: 7.75, P=0.01; red nearest islands: F=10.29, P<0.0001; blue nearest islands: F=12.26, P<0.0001). There were also interactions between island area and distance to the nearest mainland, indicating red chromaticity was higher (F=10.29, P<0.0001) and blue was lower (F=6.38, P=0.01) on more distant islands regardless of island size (Table S4). For male blue chromaticity, there was an interaction between island area and geographic region, indicating that although blue chromaticity in male passerines in the Palearctic decreased on larger islands, it increased in males in Afrotropical and Australasian regions (Table S4, F=3.03, P=0.02).

## DISCUSSION

Plumage coloration is an important phenotype involved in inter- and intraspecific communication signals, crypsis, and thermoregulation. As it is shaped by a variety of ecological and biological pressures, general patterns of colour evolution, especially in an entire order of birds, are difficult to disentangle. Our phylogenetic comparative analysis revealed plumage colouration of island species differs from their mainland counterparts. However, this pattern is more complex that has been reported previously, is mediated by a number of ecological factors, and varies across taxa. Overall, female passerines on islands exhibited reduced red and enhanced blue colouration but this effect varied among families, with some families showing significant decreases, while others increased in red and blue chromaticity between the mainland and islands. Female and male colour variation was also related to ecological covariates, including diet, latitude, habitat, and geographic region. Further, among island species, colour variation was affected by island size and isolation. Our results support the hypothesis that colour is affected by biological and ecological factors (diet, resource availability, temperature, predation and competition) as well as evolutionary history (family lineages), highlighting the complexity of colour evolution in birds.

The reduced red colour in island females suggests a reduction in carotenoid-based colouration. Carotenoid-based colouration is obtained through the consumption, metabolic conversion, and deposition of carotenoid pigments, so our observed reduction in red chromaticity may reflect variation in diet rather than an adaptation to the island environment. The reduction in red colouration could be attributed to reduced availability of carotenoid precursors in the environment or reflect increased intraspecific competition for sources rich in carotenoid precursors (54). As an example, when introduced to the Hawaiian Islands, house finches (*Carpodacus mexicanus*), which typically exhibit a red head and breast patches, became orange or yellow soon after being established and carotenoid-restricted diet experiments resulted in the loss of red plumage in male house finches (21). Alternatively, the decrease in red chromaticity may be a result of relaxed social and/or sexual selection. As islands generally exhibit lower species diversity, the reduction in sympatric species may diminish the necessity of plumage elaboration for species recognition (11). Our results also revealed an increase in female blue chromaticity on islands. If island birds are carotenoid-deficient, populations may have adapted colouration strategies by shifting endogenous precursors to melanin-based colour. One study (11) previously reported that the reduction in plumage brightness in island birds was not associated with increased black coloured plumage, such as through status signals like melanin-based badges (55),(56), but rather a continuous shift toward duller colours. This shift may be caused by increased melanin or carotenoid content in the feathers, both of which could create thicker keratin cortexes in feathers and reduce the incoherent scattering of light necessary for blue-shifted reflectance (57),(58). Whether island birds are indeed carotenoid deficient is not known; however, supplemental feeding experiments on dull island birds would be a useful study. Further research is also needed to investigate the mechanisms of reduced structural colouration and spectrometry along with microscopy of feather nanostructure to elucidate this finding.

One ecological factor that was consistently identified in our analyses as an important predictor of colour was latitude. Gloger’s rule predicts lighter coloured individuals are found at higher latitudes and darker individuals at lower latitudes (31). This rule is broadly supported in birds (59); however, the few comprehensive studies assessing latitudinal effects on colour failed to consider the consequences of island habitats or other ecological and biological explanations. Our results from phylogenetic path analyses indicate that in male Passeriformes, the direct and indirect effects of latitude on land classification are the best predictors of colour. In females, although there was an overall difference in colour between island and mainland species, this varied by latitude, where red chromaticity in island females was positively related to latitude, while mainland red chromaticity was negatively related to latitude. However, neither land classification or latitude was selected in the top path analysis model for blue chromaticity. One reason latitude may influence colour evolution is its link to temperature and precipitation, which may have direct or indirect effects on plumage coloration (60). Geographic region was the sole predictor of female blue chromaticity in our path analyses, which may be operating similarly to latitude. Habitat was another important variable identified in our PGLS models and in the path analysis for female red colour. Ambient light may vary among habitats, so selection for crypsis or conspecific signalling may vary given light environment contexts (32).

An interesting result of our study was island living influenced colour in only females. Colour elaboration in female passerines may be an adaptation to non-migratory life histories (61), which is the case for many island species (6). Although our results for red chromaticity do not support this notion, blue chromaticity increased on islands supporting the hypothesis that sedentary island life increases at least some aspects of plumage elaboration. Male colouration was not affected by island living, based on our PGLS, but rather was dependent upon other ecological factors such as diet, habitat, and latitude. In island species, island size and the distance to other islands and the mainland were significant as others have found (20), however these were not identified as important in the path analyses. Taken together, our results suggest that male colouration is influenced more by ecological and biological factors such as diet and habitat, while female coloration is affected by the combination of island living and the other ecological and biological factors.

Macroevolutionary studies are powerful ways to investigate large scale evolutionary patterns; however, they can mask differences at finer taxonomic scales (33),(62). We addressed this issue by analyzing colour evolution at the family-level. Despite finding no overall difference between mainland and island male coloration, several families showed significant increases or decreases (Fig. S2). If we had not undertaken the additional family-level analyses, we may have rejected the hypothesis that male colour differed between mainland and island species. Similarly, in females we found an overall decrease in red and an increase in blue chromaticity; however, the family-level analyses revealed the direction of these effects varied among families (Fig. S2), further highlighting how complex phenotypes, such as plumage colour can be affected by different selective forces at different taxonomic scales (63). Therefore, we join the call to urge future macroevolutionary studies to consider a range of taxonomic scales to elucidate the evolution of phenotypes that are likely being pulled in multiple directions due to differing selective pressures (33).

## Acknowledgments

We thank J. Dale for sharing the RGB colour scores (43). Funding for this project was provided by Natural Sciences and Engineering Research Council (NSERC) Discovery Grants to MWR and an NSERC Undergraduate USRA award to MDO.

---

**Table S1.** Full and reduced model selection results to assess the relationships between island and mainland classification, biological, and ecological variables on red and blue chromaticity in female and male Passeriformes birds.

| Sex | Variable | Model | K | AICc | ΔAICc | Likelihood | AICcWt | LL | Cum.Wt |
| --- | --- | --- | --- | --- | --- | --- | --- | --- | --- |
| Female | Red chromaticity | Reduced | 26 | -18362.5 | 0 | 1 | 0.72 | 9207.40 | 0.72 |
|  |  | Full | 27 | -18360.6 | 1.88<sup>1</sup> | 0.39 | 0.28 | 9207.47 | 1 |
|  | Blue chromaticity | Reduced | 24 | -15856.7 | 0 | 1 | 0.59 | 7952.50 | 0.59 |
|  |  | Full | 27 | -15856 | 0.73<sup>2</sup> | 0.69 | 0.41 | 7955.16 | 1 |
| Male | Red chromaticity | Reduced | 24 | -16349 | 0 | 1 | 0.77 | 8198.64 | 0.77 |
|  |  | Full | 27 | -16346.6 | 2.40<sup>3</sup> | 0.30 | 0.23 | 8200.47 | 1 |
|  | Blue chromaticity | Reduced | 23 | -14604.9 | 0 | 1 | 0.91 | 7325.58 | 0.91 |
|  |  | Full | 27 | -14600.3 | 4.59 | 0.10 | 0.09 | 7327.33 | 1 |

$\Delta$ AICc<4 model comparison results: <sup>1</sup>log ratio=0.14, P=0.7; <sup>2</sup>log ratio=5.33, P=0.15; <sup>3</sup>log ratio=3.66, P=0.3, indicating reduced models outperform full models. K=number of parameters, Likelihood=model likelihood, LL=log likelihood, Cum.Wt=cumulative model weight.

**Table S2.** Phylogenetic Path Analysis model selection results testing the effect of biological and ecological variables on Passeriformes island and mainland species colour. The top model appears in bold.

| | Model | k | q | C | p | CICc | ΔCICc | L | w |
| --- | --- | --- | --- | --- | --- | --- | --- | --- | --- |
| Female Red Chromaticity | **six** | **12** | **9** | **433.6** | **0** | **451.61** | **0.00** | **1** | **1** |
|  | five | 14 | 7 | 486.7 | 0 | 500.71 | 49.10 | 2.17E-11 | 2.17E-11 |
|  | seven | 12 | 9 | 519.1 | 0 | 537.17 | 85.56 | 2.64E-19 | 2.64E-19 |
|  | one | 14 | 7 | 527.9 | 0 | 541.92 | 90.31 | 2.45E-20 | 2.45E-20 |
|  | eight | 12 | 9 | 558.0 | 0 | 576.05 | 124.44 | 9.50E-28 | 9.50E-28 |
|  | two | 14 | 7 | 565.9 | 0 | 579.88 | 128.27 | 1.40E-28 | 1.40E-28 |
|  | three | 14 | 7 | 566.5 | 0 | 580.52 | 128.91 | 1.02E-28 | 1.02E-28 |
|  | four | 14 | 7 | 568.3 | 0 | 582.36 | 130.75 | 4.06E-29 | 4.06E-29 |
| Female Blue chromaticity | **five** | **14** | **7** | **492.2** | **0** | **506.19** | **0.00** | **1** | **1** |
|  | six | 12 | 9 | 524.9 | 0 | 542.92 | 36.73 | 1.06E-08 | 1.06E-08 |
|  | seven | 12 | 9 | 527.2 | 0 | 545.23 | 39.04 | 3.33E-09 | 3.33E-09 |
|  | two | 14 | 7 | 533.2 | 0 | 547.20 | 41.01 | 1.25E-09 | 1.25E-09 |
|  | one | 14 | 7 | 533.2 | 0 | 547.25 | 41.06 | 1.21E-09 | 1.21E-09 |
|  | three | 14 | 7 | 549.3 | 0 | 563.35 | 57.16 | 3.86E-13 | 3.86E-13 |
|  | four | 14 | 7 | 549.8 | 0 | 563.84 | 57.65 | 3.03E-13 | 3.03E-13 |
| Male Red Chromaticity | **five** | **12** | **9** | **381.6** | **0** | **399.64** | **0.00** | **1** | **1** |
|  | two | 14 | 7 | 466.8 | 0 | 480.82 | 81.18 | 2.35E-18 | 2.35E-18 |
|  | six | 12 | 9 | 472.6 | 0 | 490.69 | 91.05 | 1.70E-20 | 1.70E-20 |
|  | one | 14 | 7 | 483.0 | 0 | 497.02 | 97.38 | 7.15E-22 | 7.15E-22 |
|  | four | 14 | 7 | 489.7 | 0 | 503.71 | 104.07 | 2.52E-23 | 2.52E-23 |
|  | seven | 12 | 9 | 499.2 | 0 | 517.25 | 117.61 | 2.89E-26 | 2.89E-26 |
|  | eight | 14 | 7 | 503.9 | 0 | 517.96 | 118.32 | 2.02E-26 | 2.02E-26 |
|  | three | 14 | 7 | 506.4 | 0 | 520.46 | 120.82 | 5.81E-27 | 5.81E-27 |
| Male Blue Chromaticity | **five** | **12** | **9** | **468.8** | **0** | **486.85** | **0.00** | **1.00** | **0.51** |
|  | two | 14 | 7 | 474.5 | 0 | 488.52 | 1.68 | 0.43 | 0.22 |
|  | four | 14 | 7 | 475.5 | 0 | 489.49 | 2.64 | 0.27 | 0.14 |
|  | six | 12 | 9 | 471.8 | 0 | 489.80 | 2.95 | 0.23 | 0.12 |
|  | seven | 14 | 7 | 478.9 | 0 | 492.96 | 6.11 | 0.05 | 0.02 |
|  | one | 14 | 7 | 486.2 | 0 | 500.24 | 13.39 | 0.00 | 0.00 |
|  | three | 14 | 7 | 487.0 | 0 | 501.00 | 14.15 | 0.00 | 0.00 |

**Table S3.** Full and reduced model selection results to assess the relationships between classification, biological, and ecological variables on red and blue chromaticity in female and male Passeriformes island birds.

| Sex | Variable | Model | K | AICc | ΔAICc | LL | AICcWt | LL | Cum.Wt |
| --- | --- | --- | --- | --- | --- | --- | --- | --- | --- |
| Female | Red chromaticity | Reduced | 29 | -1880.72 | 0 | 1 | 1.00 | 970.51 | 1.00 |
|  |  | Full | 37 | -1868.86 | 11.86 | 0.00 | 0.00 | 973.31 | 1 |
|  | Blue chromaticity | Reduced | 27 | -1463.26 | 0 | 1 | 1.00 | 759.63 | 1.00 |
|  |  | Full | 37 | -1446.89 | 16.38 | 0.00 | 0.00 | 762.32 | 1 |
| Male | Red chromaticity | Reduced | 26 | -1665.93 | 0 | 1 | 1.00 | 859.89 | 1.00 |
|  |  | Full | 37 | -1649.28 | 16.65 | 0.00 | 0.00 | 863.52 | 1 |
|  | Blue chromaticity | Reduced | 27 | -1338.81 | 0 | 1 | 1.00 | 697.40 | 1.00 |
|  |  | Full | 37 | -1322.6 | 16.21 | 0.00 | 0.00 | 700.18 | 1 |

K=number of parameters, Likelihood=model likelihood, LL=log likelihood, Cum.Wt=cumulative model weight.

**Table S4.**
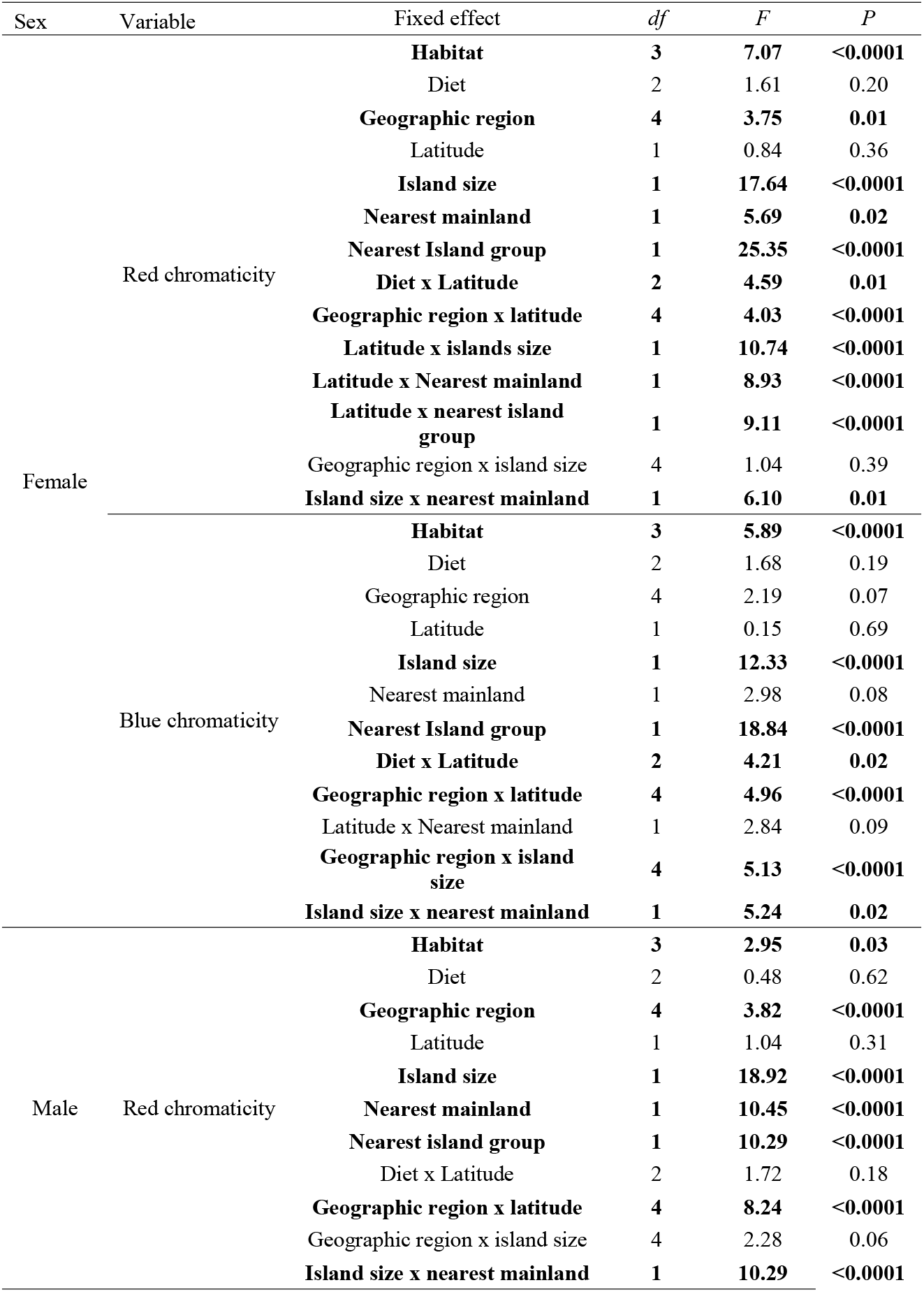
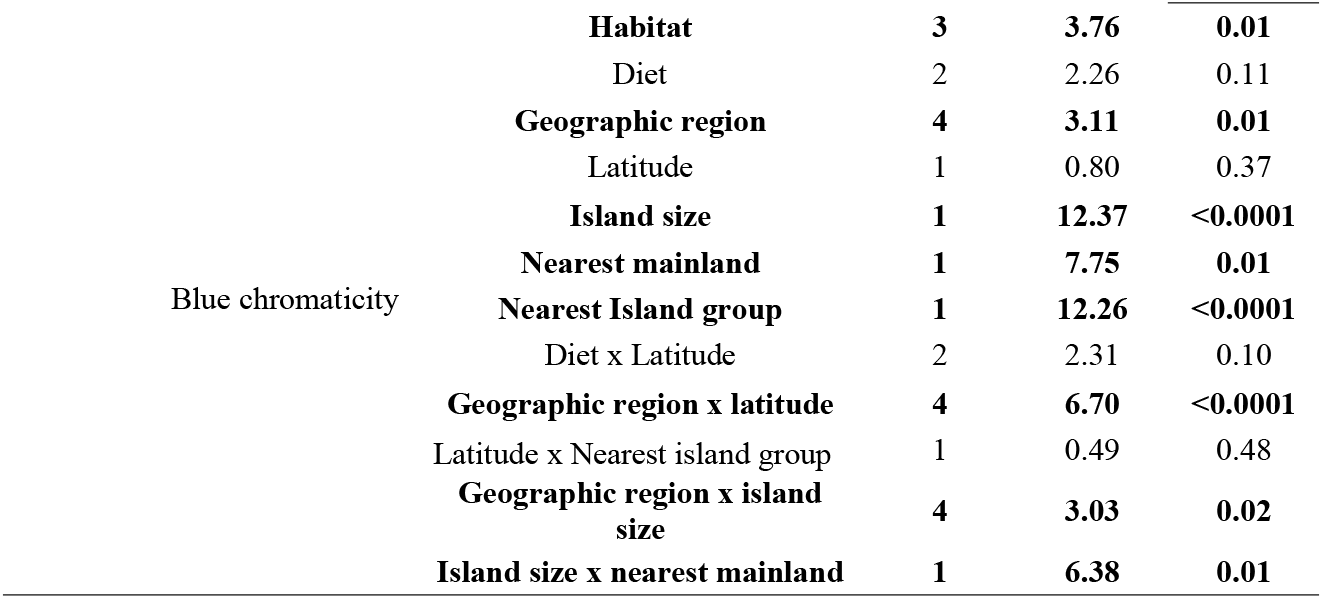
AIC selected model results demonstrating the effect of fixed effects of biological, ecological, and island characteristics on island dwelling female and male Passeriformes plumage coloration.

**Fig. S1.**
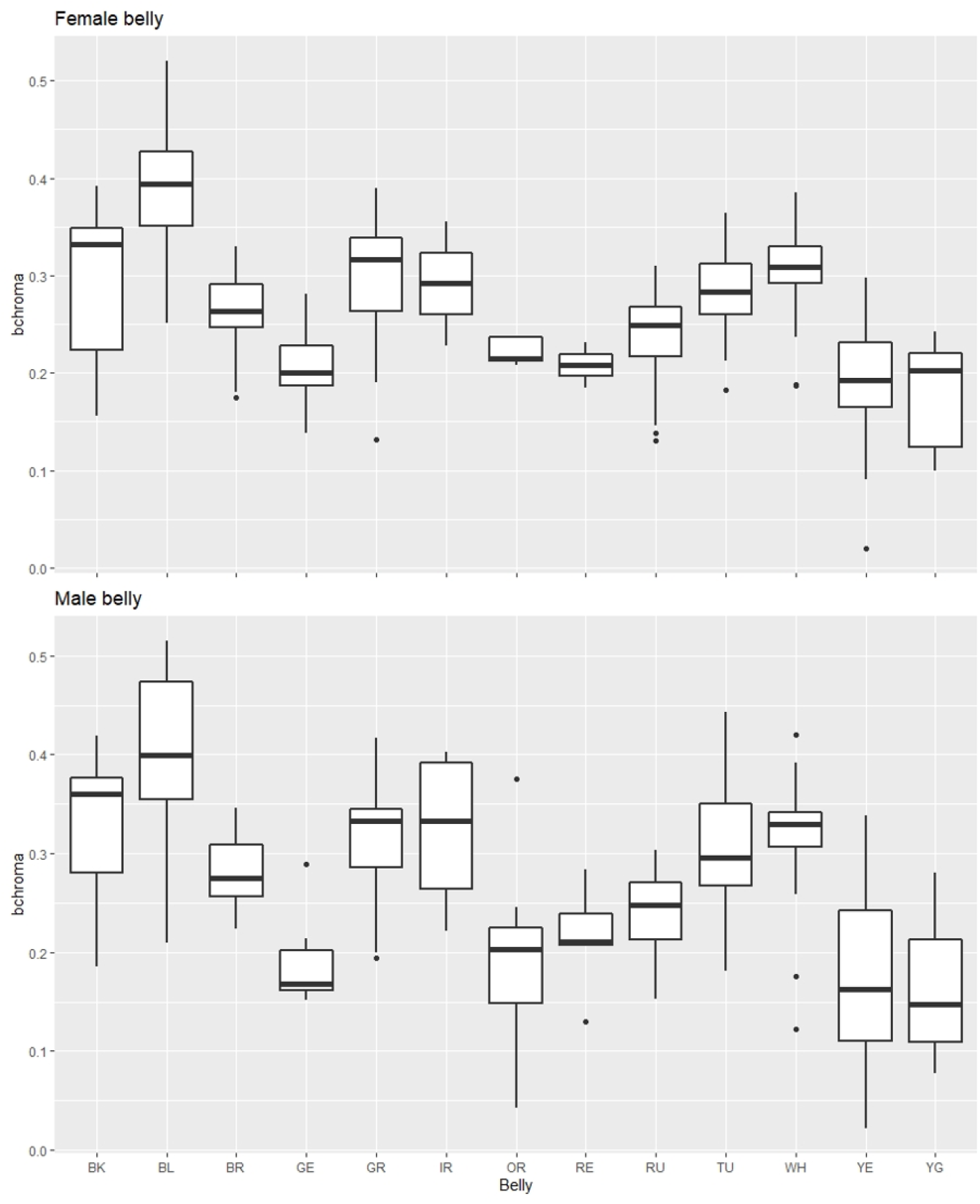
Blue chromaticity (“bchroma”, y-axis) for Thraupidae plumage patches classified as red (“RE”) or blue (“BL”) by an independent observer. There is little overlap in the distribution of chromaticity values for patches classified as red or blue, suggesting that chromaticity effectively captures the variation in red and blue plumage colouration.

**Fig. S2.**
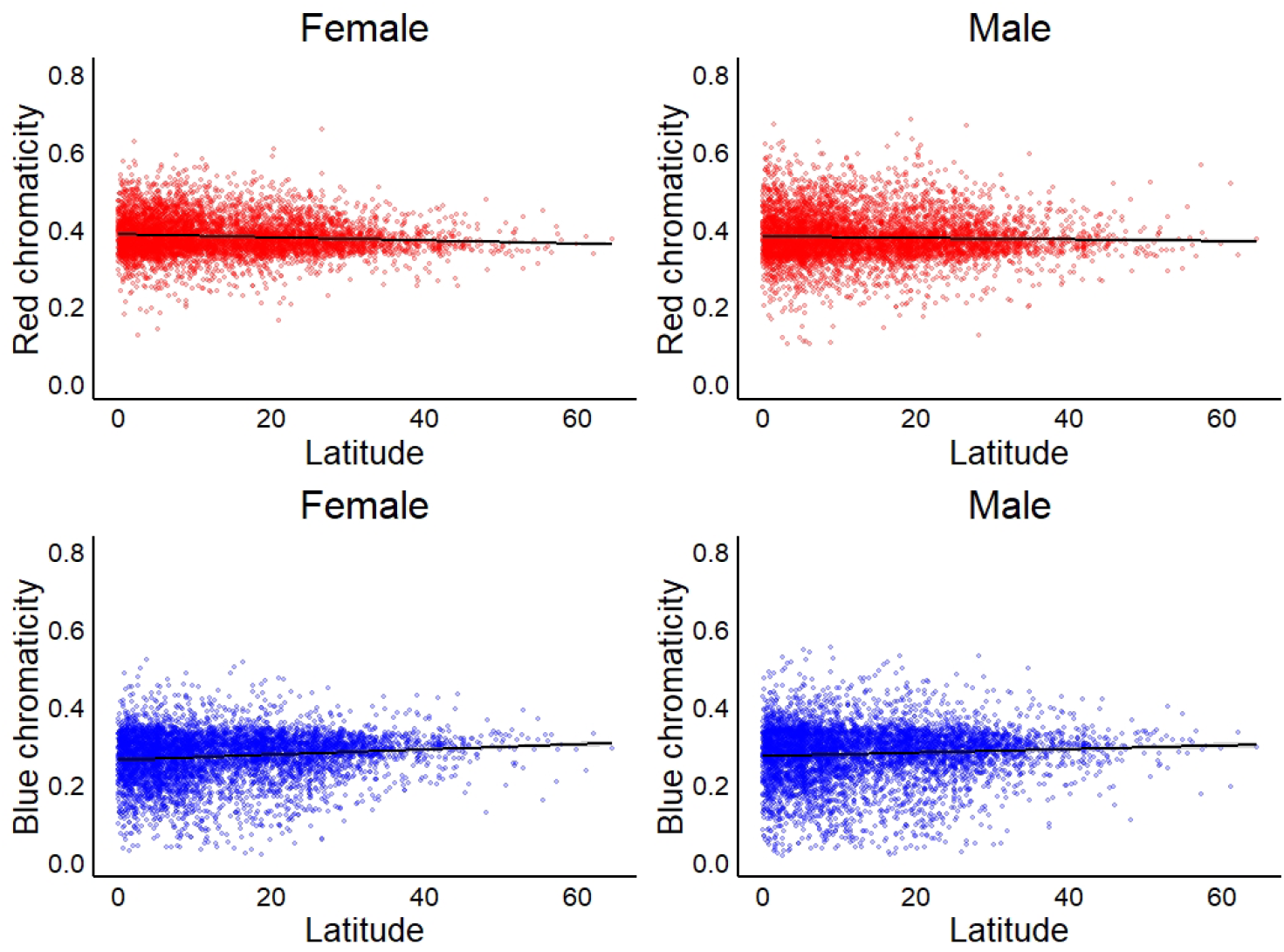
Female and male colour variation of passerine birds (n=5,693) across latitude. Top panels: Red chromaticity was significantly higher at lower latitudes for females (F=65.4, P<0.0001) and males (F=15.8, P<0.0001). Bottom panels: Blue chromaticity was positively related to latitude in females (F=75.6, P<0.0001) and males (F=14.01, P<0.0001).

**Fig. S3.**
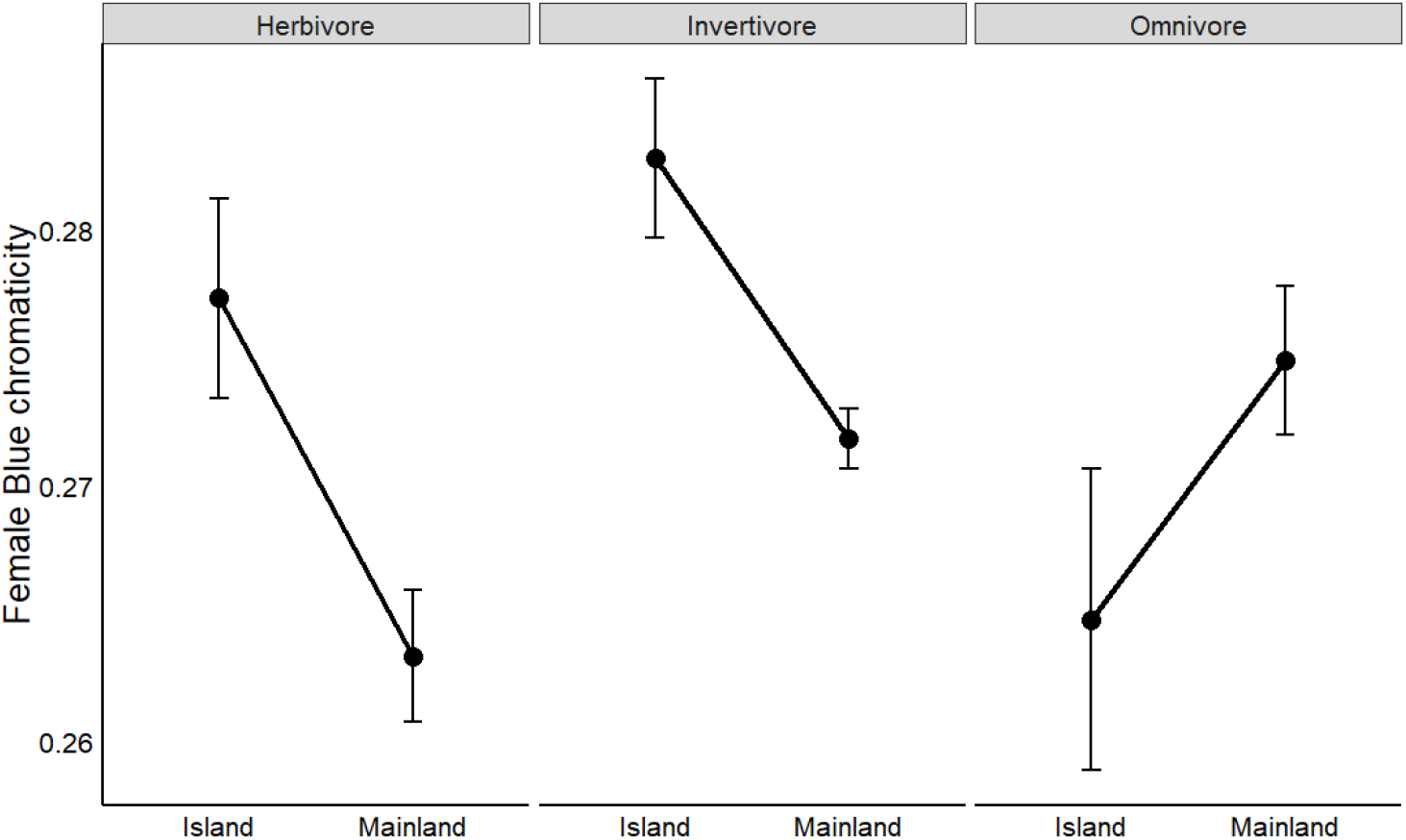
Female blue chromaticity (±SE) of passerine birds (n=5,693) of varying diets on islands and mainlands. The interaction between land type and diet indicated omnivore blue chromaticity did not vary between island and mainlands, but blue chromaticity was lower on mainland systems in herbivores and invertivores (F=17.93, P<0.0001).

**Fig. S4.**
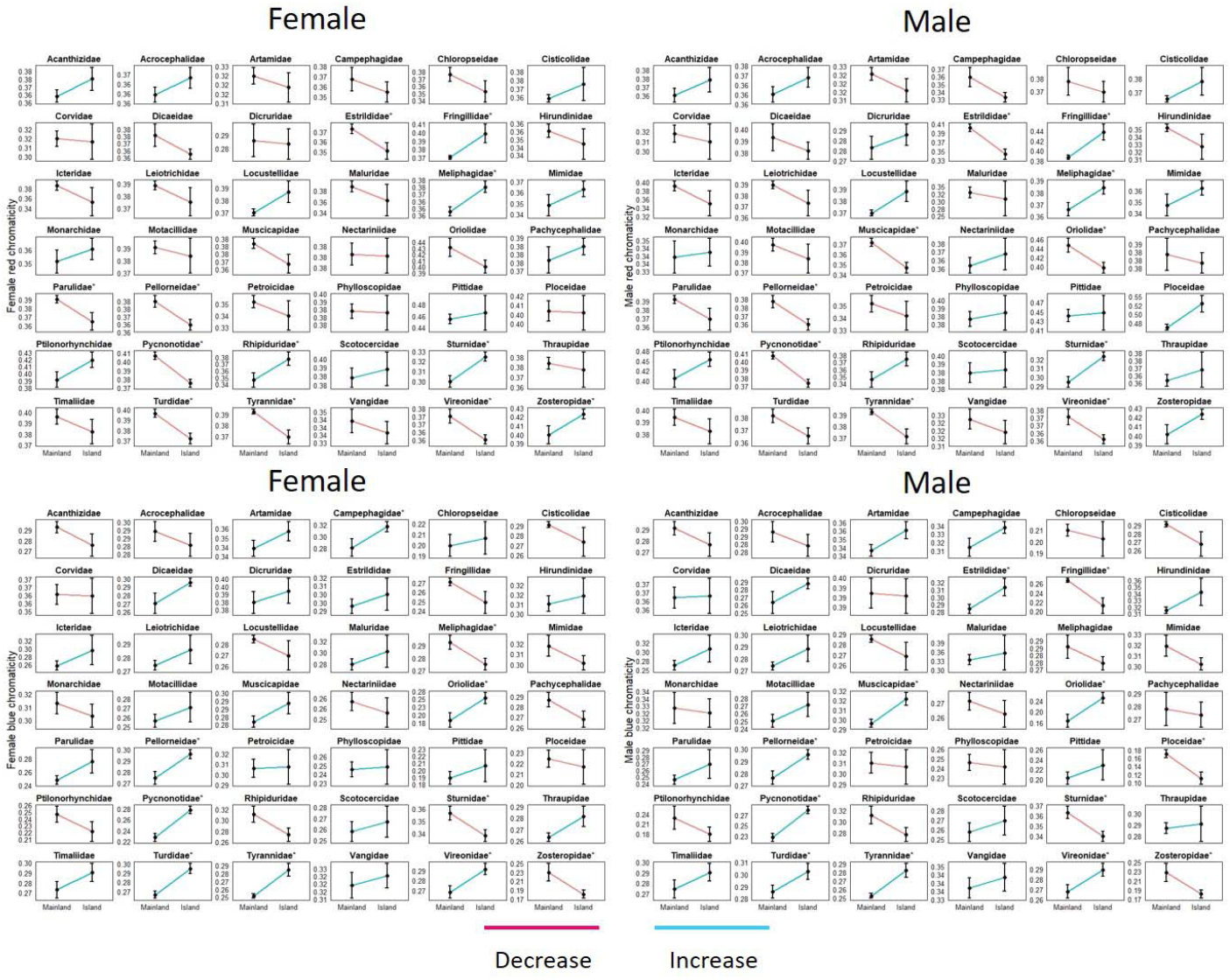
The island effect on colour evolution varied among Passeriformes families. Top panels: Red chromaticity; bottom panels: blue chromaticity. Red lines indicate decreases in chromaticity, while blue lines indicate increases in chromaticity between mainland and islands. Asterisks following family name indicate significant differences between mainlands and islands.

**Fig. S5.**
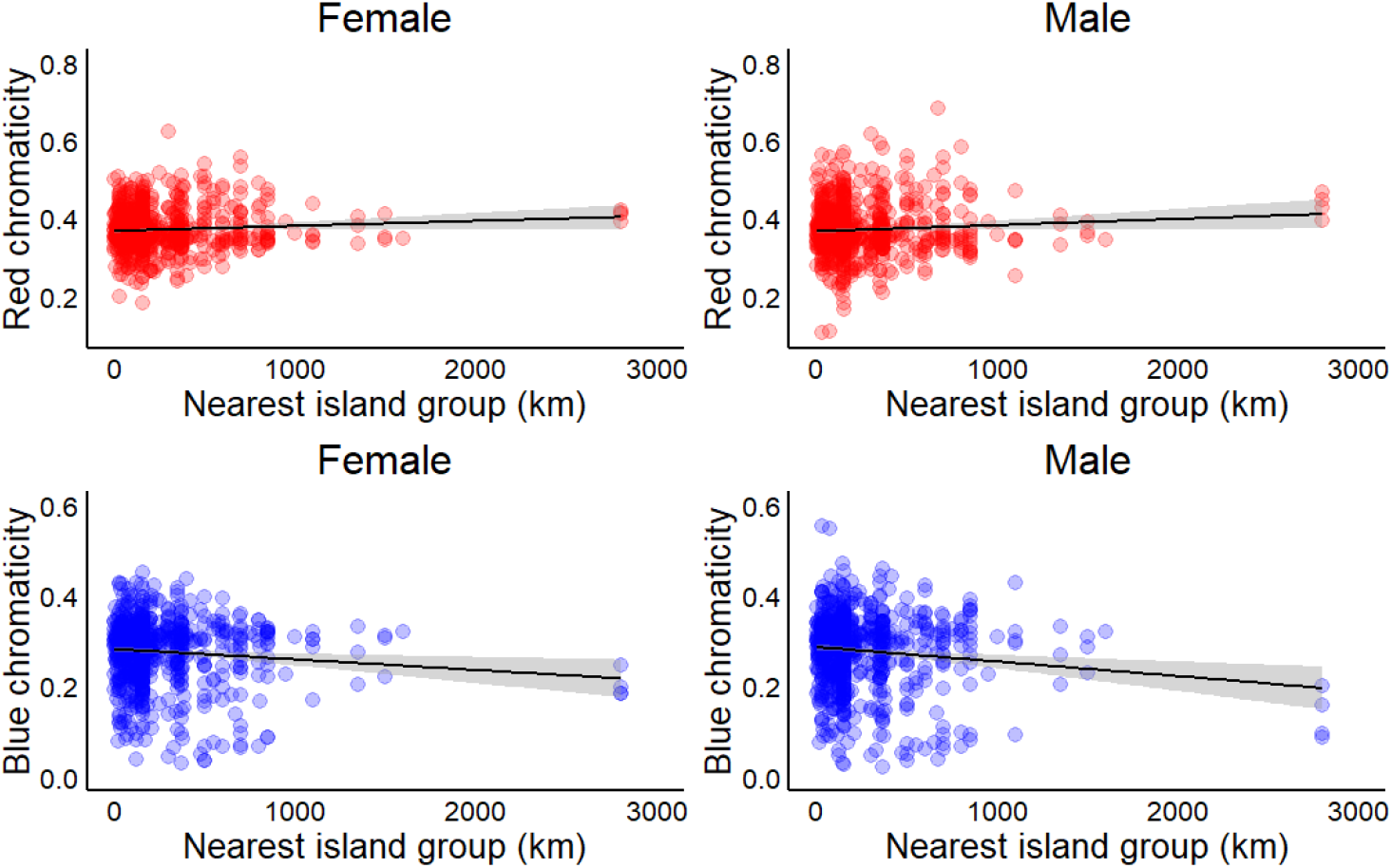
The relationship between passerine female and male chromaticity (n=1,183) and nearest island group (km). In females (F=25.4, P<0.0001) and males (F=10.3, P<0.0001), red chromaticity increased (top panels) and (bottom panels) blue chromaticity decreased (females: F=18.8, P<0.0001; males: F=12.3, P<0.0001) on more isolated islands.

